# Biophysical and Structural Characterization of a Viral Genome Packaging Motor

**DOI:** 10.1101/2022.09.25.509378

**Authors:** Nikolai S. Prokhorov, Christal R. Davis, Kashyap Maruthi, Qin Yang, Michael Woodson, Mark A. White, Lohra M. Miller, Martin F. Jarrold, Carlos E. Catalano, Marc C. Morais

## Abstract

Like many dsDNA viruses, bacteriophage λ replicates its genome as a concatemer consisting of multiple copies of covalently linked dsDNA genomes. To encapsidate a single genome within a nascent procapsid, λ must: 1) find its own dsDNA amongst the multitude of host nucleic acids; 2) identify the genomic start site; 3) cut the DNA; 4) bring the excised DNA to a procapsid; 5) translocate DNA into the capsid; 6) cut DNA again at a packaging termination site, 7) disengage from the newly filled capsid; and 8) bring the remainder of the genomic concatemer to fill another empty procapsid. These disparate genome processing tasks are carried out by a single virus-encoded enzyme complex called terminase. While it has been shown that λ terminase initially forms a tetrameric complex to cut DNA, it is not clear whether the same configuration translocates DNA. Here, we describe biophysical and initial structural characterization of a λ terminase translocation complex. Analytical ultracentrifugation (AUC) and small angle X-ray scattering (SAXS) indicate that between 4 and 5 protomeric subunits assemble a cone-shaped terminase complex with a maximum dimension of ∼230 and radius of gyration of ∼72 Å. Two-dimensional classification of cryoEM images of λ terminase are consistent with these dimensions and show that particles assume a preferred orientation in ice. The orientations appear to be end-on, as terminase rings resemble a starfish with approximate pentameric symmetry. While ∼5-fold symmetry is apparent, one of the five “arms” appears partially displaced with weaker more diffuse density in some classes, suggesting flexibility and/or partial occupancy. Charge detection mass spectrometry (**CDMS**) is consistent with a pentameric complex, with evidence that one motor subunit is weakly bound. Kinetic analysis indicates that the complex hydrolyzes ATP at a rate comparable to the rates of other phage packaging motors. Together with previously published data, these results suggest that λ terminase assembles conformationally and stoichiometrically distinct complexes to carry out different genome processing tasks. We propose a “symmetry resolution” pathway to explain how terminase transitions between these structurally and functionally distinct states.

## Introduction

Viruses are obligate intracellular parasites whose developmental pathways are initiated upon insertion of their genetic material into a host cell (1). The pathways are generally conserved within broad virus classes such as the large dsDNA viruses, including, the *Caudoviruses* (tailed bacteriophages), and the *Herpesviruses* groups (2,3). Upon insertion of their genetic material into a host cell, viral genomes are typically replicated as head-to-tail concatemers (**immature DNA**). Expression of late viral genes produces structural proteins that self-assemble into procapsid shells into which the newly replicated DNA is actively packaged. **Genome-packaging**, which is catalyzed by **terminase enzymes**, represents the intersection between the DNA replication and capsid assembly pathways (4-7); this involves processive excision of individual genomes from the concatemer and simultaneous packaging of the “mature” DNA into a preassembled procapsid shell (**Figure 1A**). Both reactions are catalyzed by the virus-encoded **terminase enzyme**, which thus performs two essential functions; (i) nucleolytic excision of individual genomes from a concatemeric precursor (**maturation reaction**) and (ii) concomitant translocation of viral DNA into the procapsid (**packaging reaction**) To accomplish these distinct tasks, terminase complexes must sequentially alternate between distinct maturation and packaging configurations during processive genome packaging^1^.

**Figure 1.**
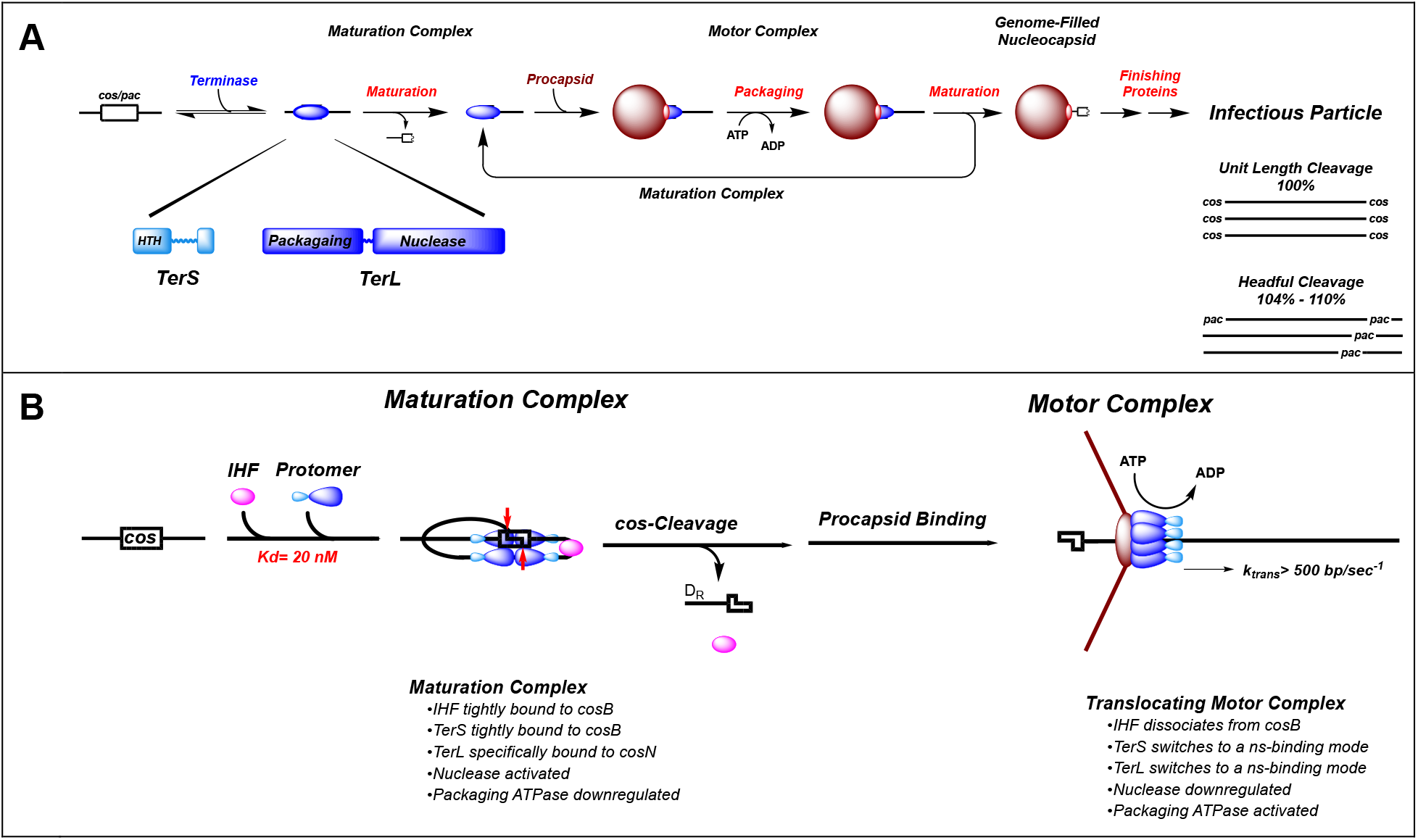
Genome Packaging in the Complex Double-Stranded DNA Viruses. <u>*PanelA*</u>. Processive Genome Packaging. Terminase enzymes are responsible for processive excision of an individual genome from a concatemeric packaging substrate (**genome maturation**) and for translocation of the duplex into a pre-formed procapsid shell (**genome packaging**). The functional enzymes are composed of large catalytic (**TerL**) and small DNA recognition (**TerS**) subunits, both of which possess conserved functional domains, as depicted at bottom, left. Two basic strategies for genome packaging, unit-length and headful, are summarized at bottom, right. Processive packaging requires that terminase cycles between a maturation complex, tightly bound at a *pac* or *cos* site in the viral genome, and a dynamic motor complex. Details are provided in the text. <u>*Panel B*</u>. The l Terminase Maturation and Motor Complexes. Four λ terminase protomers and IHF cooperatively bind to the *cos* sequence to assemble a dimer of dimers nuclease complex and position a head-to-head TerL dimer symmetrically bound to the *cosN* subsite. Assembly down-regulates the packaging ATPase site and activates nuclease activity, which introduces symmetric nicks into *cosN*, 12 bases apart (red arrows). Upon binding to the capsid portal, the head-to-head dimer of dimers nuclease complex must re-orient the protomers to engender a parallel ring-like motor complex for binding to the portal vertex. TerL and TerS subunits are depicted in blue and cyan, respectively. Details are provided in the text.

The terminase enzymes function as heterooligomeric complexes consisting of a large catalytic subunit that contains both maturation and packaging activities (**TerL**) and a small DNA recognition subunit that is required for site-specific assembly at the packaging initiation site (**TerS**; **Figure 1**) (4-7). Both subunits are essential for virus development *in vivo*. Structural studies have uncovered molecular details of isolated TerL and TerS subunits and have provided significant insight into structure-function relationships of the enzyme subunits. With the exception of bacteriophage asccphi28 (8,9), isolated TerL subunits are monomers while the TerS subunits assemble into ring-like complexes with central channels of varying stoichiometries and dimensions. This observation has led to a controversy as to whether DNA passes through the central channels or whether the complexes wrap the duplex around the TerS ring exterior (10-17); however, these interpretations are complicated by the fact that (i) the TerS structures were obtained in the absence of the cognate TerL subunit (and *vice-versa*) and (ii) they were all examined in the absence of DNA, both of which are required for the assembly of a biologically relevant complex.

Genetic and biochemical data indicate that functional enzyme complexes composed of both TerL and TerS subunits (the **holoenyzme**) are required to assemble the maturation complex at the packaging initiation site in viral DNA (*pac* or *cos*) to initiate genome packaging. Unfortunately, there is little information on the holoenzyme complexes assembled from both subunits, largely due to the challenge of recombinantly expressing and/or assembling a functional TerL:TerS complex. An exception is phage λ terminase, which is isolated as a stable heterotrimer of TerL and TerS subunits (18,19).

Bacteriophage λ is composed of a linear 48.5 kb dsDNA genome tightly packaged within a nearly icosahedral capsid shell wherein true icosahedral symmetry is broken by the presence of a tail structure situated at a unique vertex (the portal vertex) of an otherwise icosahedral shell. Typical of the complex dsDNA viruses, injection of l DNA into an *Escherichia coli* cell initiates the infection cycle, which ultimately affords linear concatemers of the λ genome and packaging-competent procapsids. Phage λ assembly has been fully reconstituted *in vitro* and an infectious virus may be assembled using purified proteins and commercially available λ DNA; this experimental accessibility allows mechanistic interrogation of each step along the virus assembly pathway. With respect to DNA packaging, we have performed detailed kinetic, biophysical and structural analyses of the maturation and packaging reactions *in vitro*. Consistent with *in vivo* data, the *cos*-cleavage endonuclease (maturation) reaction is partially stimulated by ATP/ADP, but requires the *Escherichia coli* integration host factor (**IHF**) for full activity (5). Once the genome end has been matured, the terminase motor then translocates DNA at a rate of 600 bp/sec, fueled by ATP hydrolysis, and ultimately fills the shell with DNA at liquid-crystal density which generates an internal pressure of 25 atmospheres.

As noted above, unlike most terminase enzymes, λ terminase purifies as a stable, homogeneous heterotrimer composed of two TerS subunits tightly associated with a single TerL subunit [TerL•TerS_2_, the **terminase protomer**], which is catalytically inactive (18,19). The protomer undergoes a salt-linked self-association reaction at elevated concentrations in solution to afford a multimeric ring-like **holoenzyme complex** (*K*_*D,app*_∼ 3-4 μM) that activates the catalytic activities of the enzyme (20,21). At physiological concentrations, the protomer and IHF cooperatively assemble at the *cos*-sequence of λ DNA at physiological concentrations (*K*_*D,app*_∼ 20 nM), which subsequently activates *cos*-specific nuclease activity and down-regulates packaging ATPase activity (**Figure 1B**) (20,21). Based on extensive biophysical data, we have proposed that this *maturation* complex is composed of a head-to-head dimer of dimers with approximate D2 symmetry that is thus capable of introducing symmetric nicks at the *cos*-site of the anti-parallel dsDNA, in analogy to the type IIE and IIF restriction endonucleases (**Figure 1B**) (22-24).

The post-cleavage complex must then bind to the portal vertex of a procapsid and trigger the transition to the motor complex in which nuclease activity is down-regulated and packaging ATPase activity is upregulated to power DNA packaging (**Figure 1B**). In contrast to the maturation complexes discussed above, translocating packaging motors from several systems, including phages ϕ29, λ, T4 and P74-26, among others, have been examined in great detail. Biochemical, biophysical and structural characterization of several packaging motors indicate that they function as *pentamers* of TerL subunits (5). Moreover, genetic and biochemical studies have shown that C-terminal residues in λ, P22, T3 and T4 TerL subunits interact with the portal (5), and in the ϕ29, structural studies unambiguously show that the CTD of the ATPase interact with the portal vertex (25-27). Thus, it has long been presumed that the packaging motors are composed of TerL subunits assembled in a ring-like complex, oriented in a parallel manner, as depicted in **Figure 1B**. Thus, the stoichiometry and orientation of l TerL subunits assembled on the DNA in the maturation complex vs. that assembled at the portal in the DNA translocating motor complex may be distinct, ostensibly a direct consequence of their vastly different biochemical roles in genome packaging. This raises a fundamental question: is the head-to-head, dimer-of-dimers stoichiometry of the λ maturation complex retained as it transitions to a dynamic motor complex, which would be in contrast to all other systems, or is a “fifth” protomer recruited to engender a pentameric motor.

Unfortunately, there is no structural data for a motor composed of both TerL and TerS subunits in any system. Here, a combination of analytical ultra-centrifugation (**AUC**), charge detection mass spectrometry (**CDMS**), small angle X-ray scattering (**SAXS**), enzyme kinetics and class averaging of 2D cryo-EM images to investigate the structure and function of a lambda terminase assembled from both TerS and TerL proteins. Our results suggest that the structure and stoichiometry of lambda terminase is reconfigured for maturation and translocation tasks. We propose a ‘symmetry resolution’ pathway to explain how lambda terminase transitions between these distinct structural and functional states. In addition to illuminating how quaternary reconfiguration of a molecular motor allows for multi-tasking, the ability to study functional transitions in an isolated terminase ring motors will facilitate rigorous biophysical analysis, kinetic analysis, and high-resolution structure determination, expediting a comprehensive understanding of a complex, multi-tasking molecular machine.

## Results

### Small-Angle X-ray Scattering

Understanding how terminase enzymes carry out coupled maturation and translocation activities has been limited by the inability to assemble terminase holoenzymes that include both of the required TerL and TerS subunits. An exception is phage λ terminase, which has been extensively characterized using biochemical, kinetic, biophysical and single molecule approaches (4,5,28); unfortunately, structural studies on λ terminase have been limited. As a first step towards deciphering size and shape information for λ holoenyzme, we conducted solution X-ray scattering experiments on λ protomers assembled into the catalytically competent ring complex. No radiation damage in the assembled complexes was detected over the 10– 12-hour exposure time, so the data obtained after this long exposure was used for analysis. The scattering data are not dependent on the protein concentration, as the scattering curves superimposed well for protein samples at concentrations ranging from 0.25 to 0.80 mg/mL (**Figure 2A**). This is consistent with our AUC data demonstrating that a single species, the assembled ring, predominates in this concentration range (*vide infra*).

**Figure 2.**
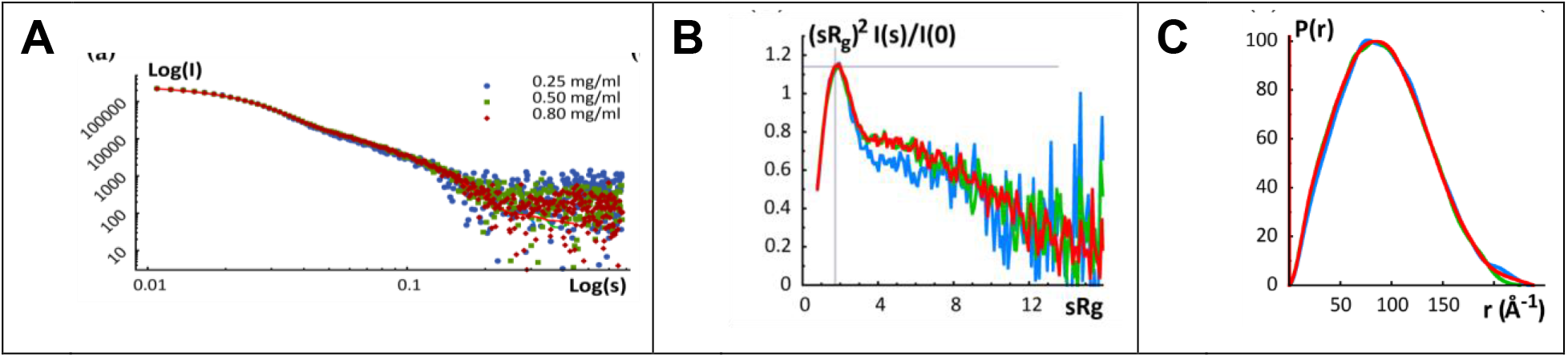
Figure 2. SAXS Analysis of the Assembled Terminase Holoenzyme. <u>*Panel A*</u>. Experimental scattering patterns for three concentrations of λ terminase scaled to concentration. Kratky plots (<u>*Panel B*</u>) and pair-distance distance distribution functions P(r) (<u>*Panel C*</u>) of the assembled λ termiase ring for the three concentrations.

Examination of the Kratky plot indicated that the complex is well-folded overall but indicated that there are likely small regions of polypeptide that are not well ordered (**Figure 2B**). Two independent methods, the Guinier approximation and the pair-distribution function P(r), were used to calculate the radius of gyration (Rg) value for each protein concentration. Both methods provided similar Rg values, with ≈ 70 ± 1 Åcalculated from the Guinier approximation and ≈ 71.9 ± 0.6 Å estimated from the pair-distance distribution function P(r) (**Table 1**). The maximum dimension of the particle, D_max_, was found to be ≈ 230 Å by examining where the pair-distance distribution function went to zero (**Figure 2C**). The Molecular weight, determined using the Porod-invariant volume method, corresponded to average molecular masses between 51.8 and 69.1 kDa, indicating that terminase forms a higher order assembly in solution consisting of ∼4.4 protomers (**Table 1**).

**Table 1.** SAXS Parameters for the Assembled Terminase Holoenzyme.

| Protein<br>Concentration | Rg<br>(Guinier)<br>(Å) | Dmax<br>(Å) | Rg (P(r))<br>(Å) | Estimated<br>Molecular Mass<br>(kDa) | Number of<br>Monomers |
| --- | --- | --- | --- | --- | --- |
| 0.80 mg/mL | 70.5(12) | 233 | 71.9(6) | 50.4 | 4.4 |
| 0.50 mg/mL | 74.1(24) | 228 | 71.1(8) | 50.0 | 4.2 |
| 0.25 mg/mL | 71.2(23) | 241 | 73(1) | 58.7 | 5.1 |

### SAXS Ab Initio Shape Calculations

The shape of the assembled terminase holoenzyme in solution can be approximated from the shapes of the Kratky plot (**Figure 2B**) and the pair-distance distribution function (P(r) (**Figure 2C**). The Kratky plot shows a bell-shaped peak at low angles that indicates a well-folded protein, and the pair-distance distribution function, P(r), shows a characteristic shape of a conical ring-like particle (29-31).

Bead models of the assembly, which approximate the molecular envelope, were then obtained using the program GASBOR. Due to the non-integer number of subunits indicated by the Porod volumes calculated at different protein concentrations (average of 4.4 protomers), GASBOR was run imposing either P4 or P5 symmetry on the resulting bead model. These molecular envelopes are indeed ring-like cylinders; models with P4 and P5 symmetry had similar dimensions: both were conical ring-like shapes with similar dimensions: the ‘base’ of the cone had a diameter of ≈ 200 - 213 Å and the height of the cones were ∼ 37 - 62 Å (**Figure 3**). Calculated scattering curves from each of the 25 envelopes with either P4 or P5 symmetry imposed fit the experimental scattering data equally well, with ∼χ^2^ values ranging from 0.82 to 0.89; models for either symmetry did not cluster within this range, and hence the ∼χ^2^ values could not be used to distinguish between potential stoichiometries.

**Figure 3.**
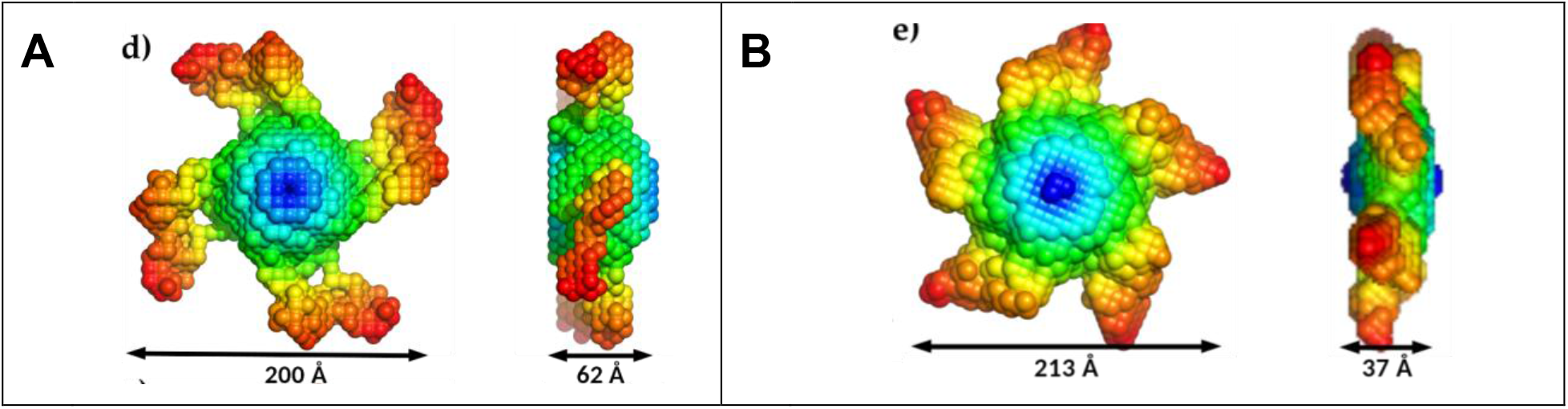
Molecular Envelopes Generated using GASBOR. Models were obtained by imposing either P4 (<u>*Panel A*</u>) or P5 (<u>*Panel B*</u>) symmetry. Both models are nearly equally consistent with the X-ray scattering curve of terminase holoenzyme.

### Hydrodynamic Modeling of the Terminase Ring

We previously employed analytical ultracentrifugation (**AUC**) to evaluate the protomer stoichiometry in the assembled λ maturation complex and we interpreted these data to indicate that the holoenzyme is composed of four protomers assembled into a ring-like structure (19,20); however, the SAXS results above are consistent with either a tetrameric or pentameric ring complex. To correlate the two studies, we modeled the hydrodynamic behavior of the structural models using the HYDROPRO program (32). The data presented in **Table 2** reveal that the predicted sedimentation coefficient of a pentameric ring (13.6 S) most closely matches the experimental sedimentation coefficient of the terminase ring previously reported (20).

**Table 2.** Hydrodynamic Modeling of the Terminase Holoenzyme SAXS Data.

|  | **Tetramer Ring** | **Pentamer Ring** |
| --- | --- | --- |
| Molecular Weight (Calculated) | 460 kDa | 575 kDa |
| Radius of Gyration | $7.4 \times 10^{-7}$ cm | $7.3 \times 10^{-7}$ cm |
| Volume | $6.5 \times 10^{-19}$ cm<sup>3</sup> | $7.4 \times 10^{-19}$ cm<sup>3</sup> |
| $S_{(20,w)}$ | 11.8 S | 13.6 S |

### Biophysical Characterization of Terminase Holoenzyme

In contrast to our published studies that suggest terminase holoenzyme is composed of four protomers both in solution (19,20) and on viral DNA (21), hydrodynamic modeling of the SAXS data above is most consistent with a pentameric structural model. Given that the protomer stoichiometry has important mechanistic implications in both the maturation and motor complexes, we re-examined the protomer stoichiometry in the assembled holoenzyme in solution using an expanded AUC approach.

We first employed sedimentation velocity analytical ultracentrifugation (**SV-AUC**) as described in Materials and Methods. Three different concentrations of protomer (2.67 μM, 4 μM and 8 μM) were examined and the data were analyzed using three independent approaches. First, the model independent time derivative approach (DCDT^+^) was employed, and the data are consistent with a single apparent species (data not shown). Analysis of the g*(s) distributions affords a *s(*_*20,w*_) of 13.1 to 13.5 S for the three protein concentrations (**Table 3**). We next used Sedfit, a direct boundary fitting approach, and the c(s) distributions are shown in **Figure 4A**. Analysis of the data yields *s(*_*20,w*_) of 13.2 to 13.3 S for the three protein concentrations (**Table 3**). Finally, we employed SEDANAL to simultaneously fit all three data sets according to a single species model. The results of the global fit are shown as solid lines in **Figure 4B**, which yield *s(*_*20,w*_)= 13.6 S (**Table 2**). In sum, the three analytical approaches afford similar values for *s(*_*20,w*_) that are commensurate with both hydrodynamic modeling of the SAXS data (**Table 2**) and our published data (19-21); however, and despite the fact that all three programs afford similar values for *s(*_*20,w*_), the derived molecular weights of the assembled holoenzyme differ between the different analytical approaches and the calculated protomer stoichiometry thus varies between 3.3 to 4.4 (**Table 3**).

**Figure 4.**
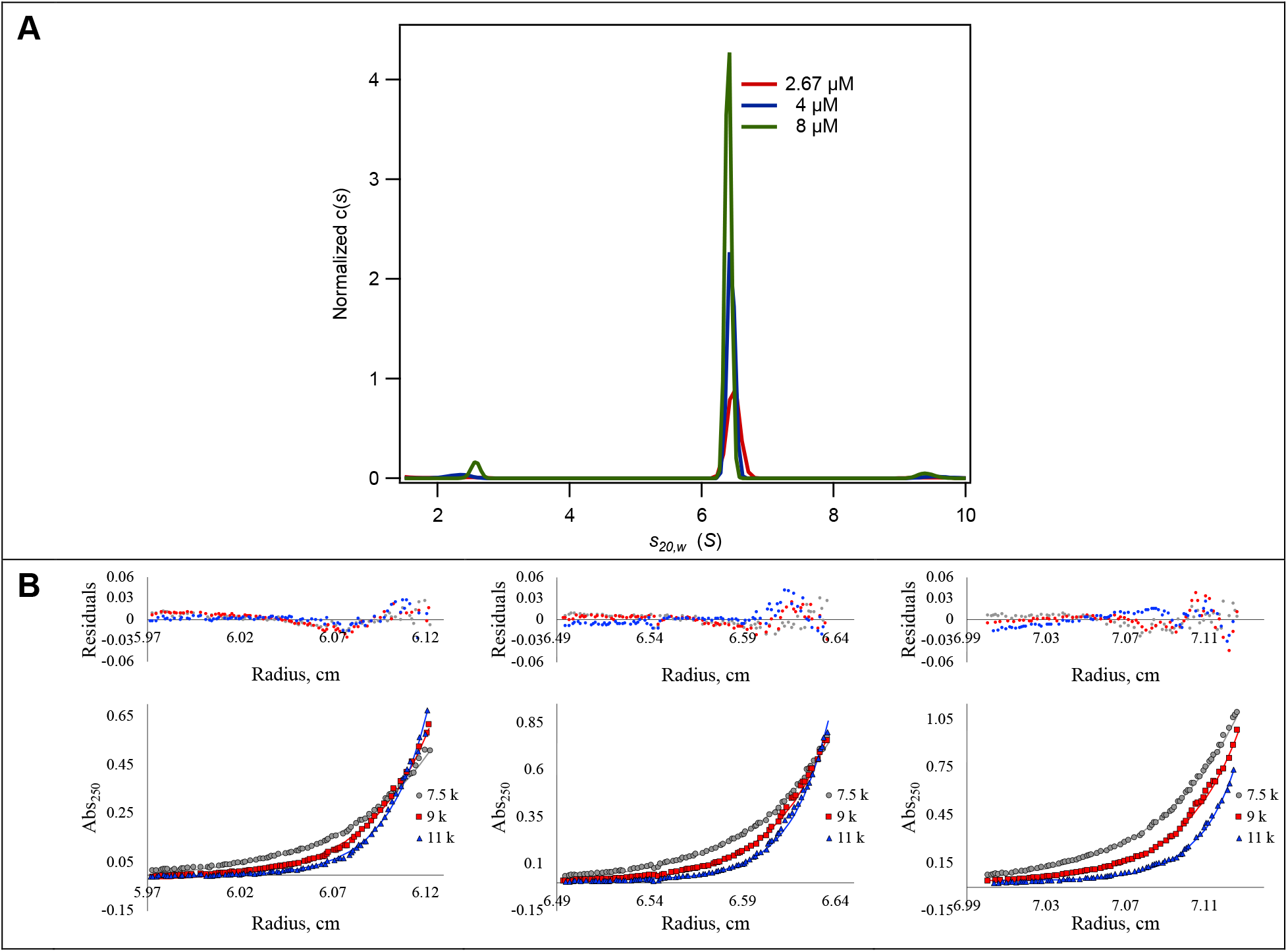
Analytical Ultracentrifugation of the Assembled Terminase Holoenzyme. <u>*Panel A*</u>. Normalized c(S(20,w)) of the SV-AUC data analyzed by Sedfit. Note the appearance of a minor 5.1 S species in all samples, which represents a small fraction (<5%) of the ring that has dissociate to the protomer. <u>*Panel B*</u>. Sedimentation equilibrium analysis of the assembled terminase holoenzyme. Sample concentrations were 2.67 μM, 4 μM and 8 μM (*left to right*) and were spun at the indicated speeds as described in Materials and Methods.

**Table 3.** AUC Analysis of the Assembled Terminase Holoenzyme Ring.

| **SV-AUC** | **Speed (rpm)** | **Method** | **$S_{(20,w)}$** | **Derived MW (kDa)** | **Derived Stoichiometry\*** |
| --- | --- | --- | --- | --- | --- |
| 2.67 mM | 32 k | DCDT+ | 13.5 | 503 | 4.4 |
| 4 mM |  | DCDT+ | 13.4 | 477 | 3.9 |
| 8 mM |  | DCDT+ | 13.1 | 481 | 3.6 |
| 2.67 mM |  | SEDFIT | 13.3 | 446 | 4.1 |
| 4 mM |  | SEDFIT | 13.3 | 437 | 3.8 |
| 8 mM |  | SEDFIT | 13.2 | 471 | 3.6 |
| 2.67 mM |  | SEDANAL | 13.6 | 416 | 4.2 |
| 4 mM |  | SEDANAL | 13.6 | 411 | 4.1 |
| 8 mM |  | SEDANAL | 13.6 | 375 | 3.3 |
| Global |  | SEDANAL | 13.6 | 384 | 3.3 |
| **SE-AUC** |  |  |  |  |  |
| 2.67 mM | 7.5, 9, 11k |  | N/A | 538.3 | 4.7 |
| 4 mM | 7.5, 9, 11k |  | N/A | 503.2 | 4.4 |
| 8 mM | 7.5, 9, 11k |  | N/A | 509.8 | 4.4 |
| Global | 7.5, 9, 11k |  | N/A | 513.5 | 4.5 |

Therefore, we next employed sedimentation equilibrium analytical ultracentrifugation (**SE-AUC**) to obtain the molecular weight of the assembled terminase holoenzyme that is uncluttered by shape information inherent in an SV-AUC experiment. The SE-AUC experiment used the same three protein concentrations employed in the SV-AUC studies (2.76 M, 4 M and 8 M) and the samples were spun at 7500 RPM, 9000 RPM and 11000 RPM as described in Materials and Methods. Based on the SV-AUC results (**Figure 4A**), the data were fit according to a single species model and a global fit of all 9 data sets (three terminase concentrations and three rotor speeds) was performed; the results are presented as solid lines superimposed on the data (**Figure 4C**). This analysis yields a derived molecular weight of 513.5 kDa which corresponds to a protomer stoichiometry of 4.5 S (**Table 3**).

### Characterization of the Assembled Ring by Mass Spectrometry

Charge Detection Mass Spectrometry (**CDMS**) was also employed to characterize the terminase protomer and assembled ring species, as described in Materials and Methods. The data for the protomer clearly demonstrates the presence of a prominent species with a molecular weight 115.3 kDa (**Figure 5A**); this is consistent with the molecular weight calculated from the gene sequence (114.97 kDa) and that obtained experimentally from AUC (115 ± 3 kDa) (19). CDMS analysis of the assembled ring species is shown in **Figure 5B** and similarly reveals a predominant species with average molecular weight 576.3 kDa; this is consistent with a ring composed of 5.0 protomers. Of note, minor species with MW= 75 kDa and 505 kDa are also observed, which likely derive from the ring pentamer that has lost one TerL subunit (74.13 kDa, by gene sequence). This is discussed further below.

**Figure 5.**
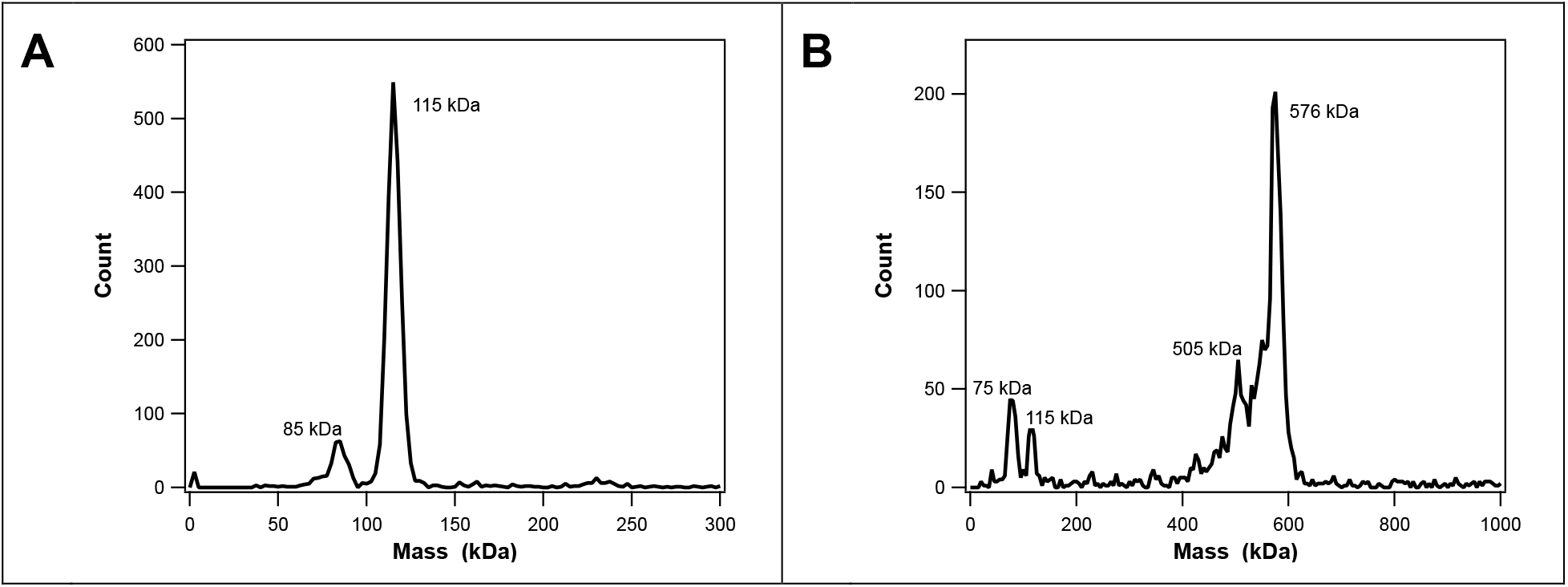
Charge Detection Mass Spectrometry (CDMS). <u>*Panel A*</u>. CDMS Spectra of the Terminase Protomer. A bun size of 5 kDa was used for the mass distribution. Gaussian fitting of the major peak gives a molecular weight of 115.434 kDa. The peaks at 85 kDa and 230 kDa likely represent the TerL subunit and a dimer of protomers, respectively. <u>*Panel B*</u>. CDMS Spectra of the Terminse Ring Species. A bun size of 5 kDa was used for the mass distribution. Gaussian fitting of the major peak gives a molecular weight of 576.332 kDa. The peaks at 75 kDa and 505 kDa likely represent an isolated TerL subunit that has dissociated from the pentamer ring, respectively. The peaks at 115 kDa and 1130 kDa likely represent free protomer and a dimer of pentamer rings.**A**

### Cryo-electron Microscopy and Image Processing

CryoEM images of terminase ring show a uniform collection of isometric particles with a radius of ∼180 Å (**Figure 6**). Most particles appear to be end-on views of ring-like particles, with occasional longer, potentially side views with approximate dimensions of 130 Å long and ∼190 Å at the widest point. The observed dimensions of side and top views are mostly consistent with the SAXS data and generated envelopes. Two-dimensional class averages of terminase complexes confirmed preferred orientations, presumably corresponding to end-on views of the terminase ring. Due to the preponderance of end-on views, the paucity of particles in other orientations, and the reduced dimensionality of particle projections observed in cryoEM, it was not possible to reconstruct a 3D volume; essentially, reliable information regarding the dimension corresponding to cone height was missing. Nonetheless, the class averages of the end-on views were very good and provide meaningful information. Rather than forming a continuous cone, terminase ring more closely resembles a conical starfish-like structure, wherein the apex of the cone corresponds to the central disk of a starfish (**Figure 6**). Further, the relative brightness of regions just outside the dark central relative to regions at more distant radial locations indicates more density is projected in the central area along the viewing directions, consistent with a cone-like structure. Importantly, like a starfish, the class averages show five radial extensions corresponding to starfish legs. It is worth noting that while the particles are clearly pentameric, some classes clearly deviate from 5-fold symmetry. In these classes one or two legs are not necessarily related to their neighbors by a strict 72-degree rotation, and sometimes appear shortened/differently oriented, with weaker more diffuse electron density. Finally, the class averages clearly show a central pore, confirming that these higher order terminase assemblies are ring-like structures, consistent with other phage terminases and ASCE ring ATPase motors (33).

**Figure 6.**
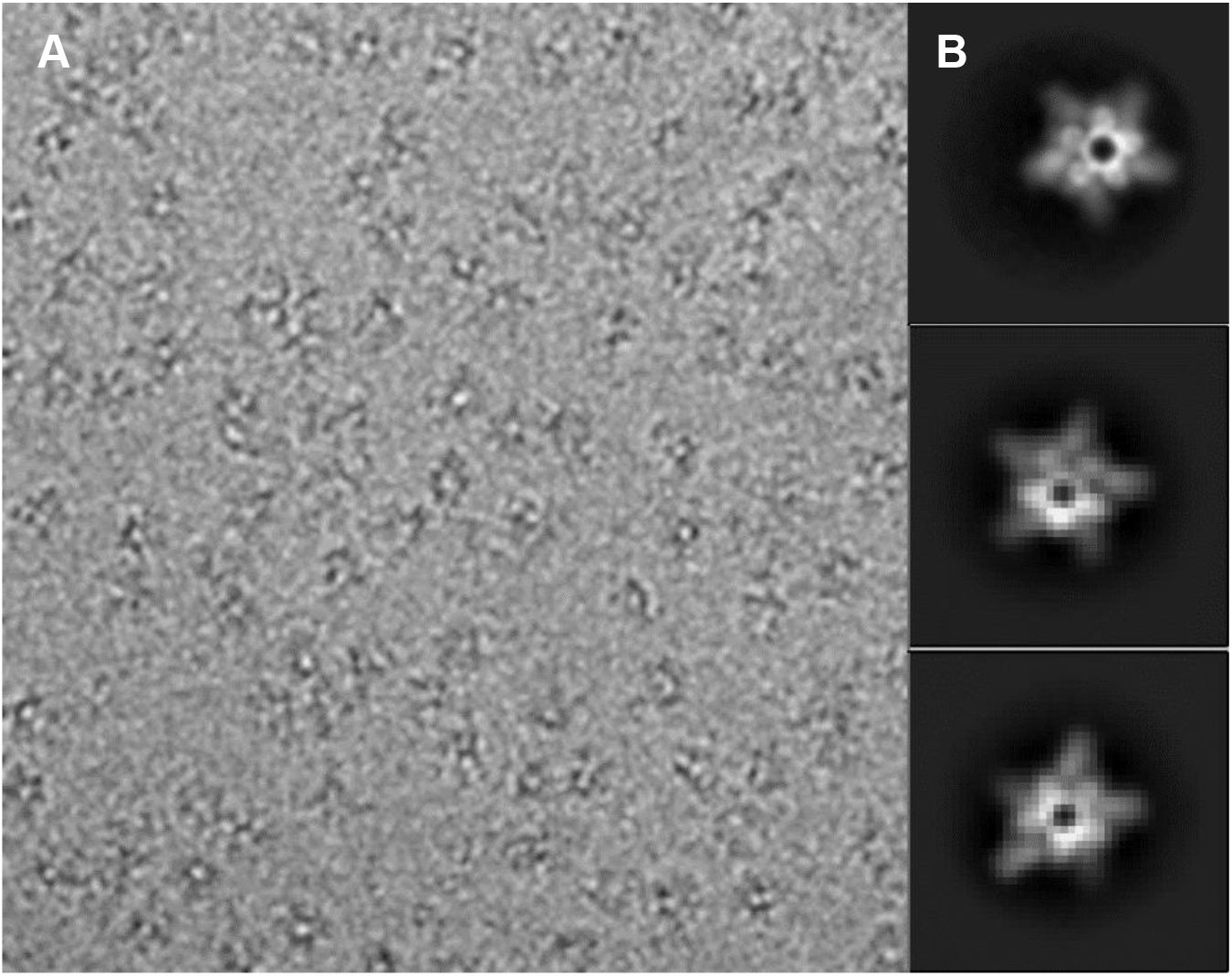
CryoEM of terminase complex. <u>*Panel A*</u>. Typical cryoEM micrograph of the terminase translocation complex showing mostly end-on views of particles. Magnification is 60,000×. <u>*Panel A*</u>. Class averages of imaged terminase particles. The relative contrast is reversed in panels A and B. Note that density corresponding to one of the five arms is weaker in the lower two panels.

### ATPase Activity

Here we describe a kinetic analysis of lambda terminase ATP hydrolysis activity. As previously demonstrated, the terminase protomer is devoid of ATPase activity, while the assembled ring efficiently hydrolyzes ATP to ADP (**Figure 7**, *inset*). We analyzed the kinetic data according to a Michaelis-Menten model, which yielded *K*_*M*_= 0.495 ± 0.070 μM, *V*_*M*_= 0.157 ± 0.006 μM/min and *k*_*cat*_= 15.7 ± 0.6 min^-1^. We note that the *K*_*M*_ obtained here is an order of magnitude lower than that previously reported (34,35).

**Figure 7.**
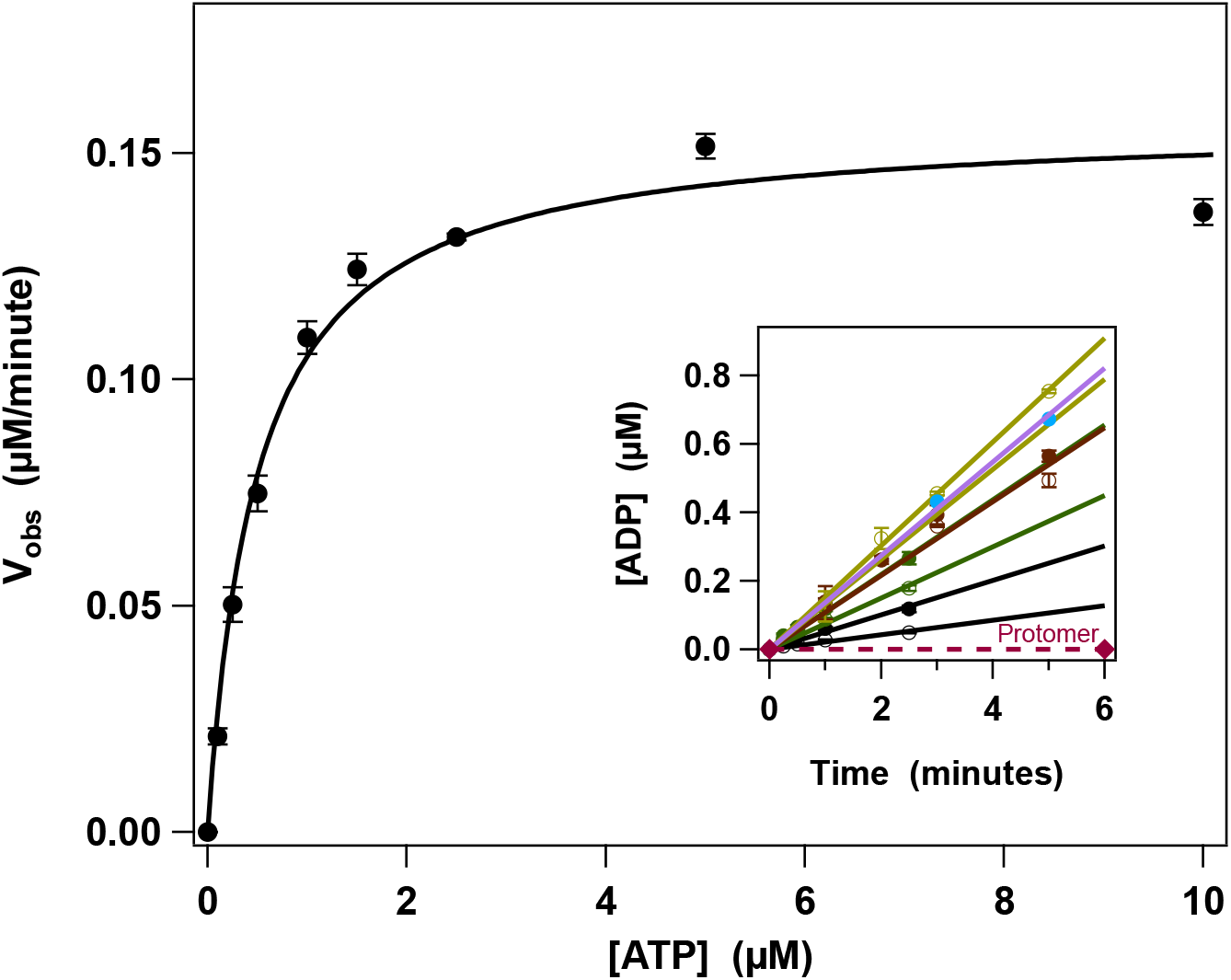
Kinetic Analysis of the Terminase Protomer and the Assembled Holoenzyme. *Inset:* Time course for ADP formation. Dashed line represents protomer data while solid lines represent data for the pentamer ring. Only the linear portions of the data were used to calculate the observed rates. Main figure shows a Michaelis-Menten plot; solid line represents the best fit of the data.

## Discussion

Genome packaging in the dsDNA viruses is highly conserved; a single genome is excised from a concatemeric precursor and concomitantly inserted into a pre-assembled procapsid (**Figure 1A**). These reactions are catalyzed by a terminase enzyme complex whose macromolecular components assemble into two distinct quaternary configurations ∼ the maturation and translocation complexes (**Figure 1B**). Transitions between these assemblies must be tightly coordinated to processively excise and package genomes from the concatemer. In bacteriophage lambda, biophysical data indicate that the conserved TerL and TerS subunits assemble into a heterotrimeric protomer (TerL•TerS_2_) which further assemble into a dimer-of-dimers maturation complex arranged with ∼ D2 symmetry. While such an arrangement is reminiscent of the type IIE nucleases, and thus consistent with the genome cutting functions of terminase, it would seem to be inconsistent with the DNA translocation (motor) function of the enzyme. Lambda terminase belongs to the Additional Strand, Catalytic “E” (**ASCE**) family of ATPase wherein higher order ASCE ATPase complexes (helicases, proteases, AAA+ proteins) are typically arranged as 5- or 6-membered rings, wherein all subunits have the same polarity and are related by a simple rotation around the central axis of the ring (5,7,33). Indeed, terminase motors characterized in other phage systems adopt this arrangement in pentameric motor complexes. Importantly, however, there is no structural information on a terminase complex assembled from both the requisite TerL and TerS subunits; as a first step in characterizing a catalytically competent complex assembled from all the necessary components, we sought to characterize the stoichiometry and symmetry of the λ terminase holoenzyme in solution.

Sedimentation equilibrium AUC analysis showed that λ terminase holoenzyme is a single species in solution, with a molecular weight of ∼514 kDa. This corresponds to a complex composed of ∼ 4.5 terminase protomers, on average. The Porod volume of the complex, estimated via small angle X-ray scattering, is similarly consistent with a complex made of ∼4.4 to 5 protomers. Additional analysis of low angle scattering data showed that protomers in the assembly were compact and mostly well-folded. SAXS data further provided information regarding the shape of the holoenzyme, suggesting the complex formed a conical assembly, with a radius of gyration of 72 Å and a maximum diameter of 230 Å. Extensive classification of 2D cryoEM images indicated that the particles adopt preferred, end-on views of the conical assembly in vitreous ice. These class averages resembled a starfish, with five extensions radiating from a central disk. Note that unlike a starfish, there was reduced density in the center of the disk, indicating a central channel and consistent with a ring-like structure. Careful examination of the 2D class averages indicated that while the radial extensions were always related by an ∼72°degree rotation, in some class averages, density for one or more of the radial extensions was weaker/diffuse and/or slightly differently positioned. Hence, the simplest explanation consistent with all the data is that the terminase ring structure is a pentamer in solution, but that the occupancy of one protomeric subunit is incomplete and/or that one subunit is arranged differently than the others.

Incomplete occupancy at one position of the terminase ring would explain the non-integer number of subunits observed in AUC and SAXS experiments, since these techniques reflect the average mass of the ensemble. Based on the known stoichiometry of the protomer and structures of other phage TerS complexes, we suspect the overall conical structure of the complex is organized by ten small TerS subunits, with five TerLs arranged around the wide end of the cone. Regarding the observed partial occupancy, we cannot be certain if the assembly is missing an entire protomer (TerL•TerS_2_) or if one protomeric subunit is missing the large TerL subunit; we suspect the latter. Indeed, CDMS data are consistent with such an interpretation. Importantly, the terminase holoenzyme investigated here hydrolyzes ATP at rates comparable to other phage terminases assembled on procapsids, consistent with a functional translocation complex.

The aggregate data presented here suggests that the λ terminase motor complex functions as a pentameric ring-like assembly in a parallel orientation with the TerL C-terminal ASCE ATPase domains bound to the portal vertex, consistent with the known structures of other phage terminases. In contrast and based on our previously published biochemical and biophysical data, we have proposed that the λ terminase maturation complex is composed of a head-to-head dimer of dimers complex assembled at *cos* (4,5,36). This model is supported with biological precedence; while tetrameric nuclease complexes are common, there are to our knowledge no examples of a pentameric endonuclease complex. Further, a parallel arrangement of subunits would seem to preclude the ability of the maturation complex to introduce symmetric nicks into the pseudo-palindromic *cosN* sequence. This begs the questions as to how terminase tranforms from the ∼ D2 tetrameric maturation complex into the cyclically symmetric pentameric ring-like translocation complex.

To explain how this reconfiguration of subunit polarity and stoichiometry occurs, we propose the following ‘symmetry resolution’ mechanism (**Figure 8**). After nicking the dsDNA duplex, the previously reported strand-separation activity of the maturation complex displaces the upstream fragment of the duplex, resulting in the loss of DNA binding energy by the upstream protomers. This disruption in turn allows a dimer of protomers to flip into a parallel orientation such that all of the TerL subunits bind to the newly formed end of DNA. This quaternary reconfiguration is driven by cooperative binding of the TerL subunits to the matured DNA end, as well as the energy released via TerS self-association interactions. As shown in the 2D class averages of the terminase translocation complex, the radial extensions presumably corresponding to individual protomers are related by an ∼72° rotation; hence four TerL subunits could not close into a ring, leaving space for fifth subunit to engage the duplex. This reorganization would arrange the C-terminal procapsid-binding residues of TerL in a parallel orientation poised to interact with the portal vertex at the unique pentameric vertex of the procapsid. Such an arrangement would result in a TerL ring-like structure that encircles dsDNA, and which is consistent with known structures of TerL translocation complexes and typical for the larger ancient and ubiquitous ASCE superfamily to which TerL belongs. The decameric TerS ring acts as a “sliding clamp” to ensure a highly processive machine, in analogy to sliding clamps involved in DNA replication (37).

**Figure 8.**
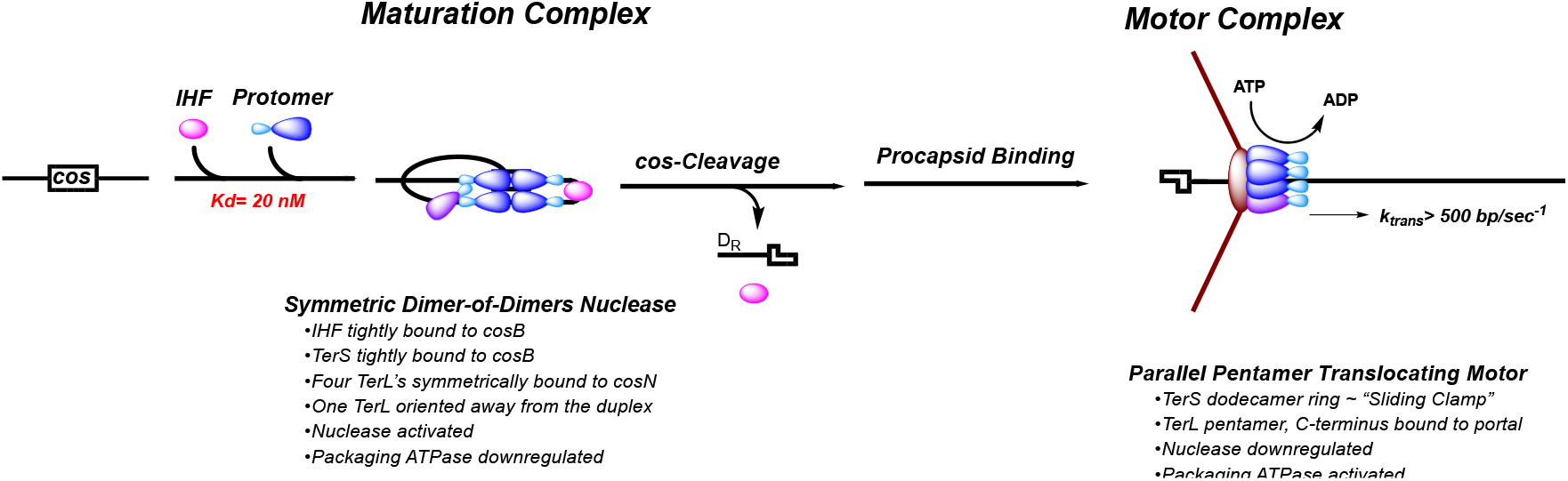
Model for the Transition to Packaging. Details provided in the text.

## Conclusions

Viral dsDNA packaging motors have been studied extensively, with a focus on understanding the mechanochemistry of DNA translocation. However, the virally encoded terminase enzymes that carry out encapsidation are versatile multi-tasking machines responsible for two related but distinct functions. Previous efforts have extensively characterized initial formation of λ terminase protomers as well as subsequent assembly of four protomers at *cos*, the packaging initiation site in λ DNA. Based on sequence similarity and common function with typ IIE nucleases, it was proposed that these four protomers are arranged with ∼ D2 symmetry at the genomic *cos* site. While such an arrangement makes sense for the nucleolytic function, it is inconsistent with the symmetry and stoichiometry of the translocation function, which is carried out by the N-terminal ASCE ATPase domain. Indeed, we show that a terminase assembly that likely reflects the translocation complex consists of five protomer subunits, consistent with the pentameric assemblies observed for other phage terminases. Our data also suggest partial occupancy for at least one of the subunits, explaining how AUC experiments performed both here and previously, along with SAXS experiments presented here, can suggest a non-integer number of protomer subunits. Hence, the λ terminase motor is capable of self-assembling substantially different quaternary configurations appropriate for distinct biophysical tasks. To better understand this functional flexibility, we propose a sequence of specific molecular events in phage λ, a symmetry resolution mechanism, that drive the site-specific assembly and activation of a stable maturation complex and its transition to a dynamic DNA translocating motor complex.

## Methods

### General

The terminase protomer and ring species were purified by published procedures 36,38). All materials were of the highest quality commercially available.

### SAXS

SAXS experiments were performed at the Sealy Center for Structural and Computational Biology (SCSB) at the University of Texas Medical Branch (UTMB). The X-ray/SAXS station at the SCSB consists of a high-brilliance FR-E++DW Superbright x-ray generator, with the industry standard RAXIS-IV++ crystallography system with both Cu and Cr optics. In addition, the source is connected to a Rigaku BioSAXS-1000 with a Kratky camera and a 96-well automatic sample changer. Samples were illuminated with X-rays generated by the Cu anode, corresponding to wavelength of 1.5418 Å. Scattering intensities I(q) for the protein and buffer samples were recorded as a function of scatter-ing vector q (q = 4πsinθ/λ, where 2θ is the scattering angle and λ is the X-ray wavelength). The sample-to-detector distance was 0.476 m, which resulted in a q range of 0.009–0.68 Å ^−1^, and all experiments were performed at 5 °C. The data collection strategy described by Hura was used in this study (39). Briefly, SAXS data were collected for three protein concentrations (0.80, 0.50, and 0.25 mg/mL) and for three matching buffer samples. For each sample measurement, SAXS data were collected with 1-hour sub-frames to assess radiation damage. The buffer scattering contributions were subtracted from the sample scattering data using the SAXNS_ES web server (https://xray.utmb.edu/SAXNS/). Data analysis was performed using the program package PRIMUS from the ATSAS suite (40,41). Experimental SAXS data obtained for different protein concentrations were analyzed for aggregation and folding state using Guinier and Kratky plots, respectively. The forward scattering intensity I(0) and the radius of gyration Rg were evaluated using the Guinier approximation: I(q) ≈ I(0) exp(−q2Rg)2/3, with the limits qRg < 1.5. These parameters were also determined from the pair-distance distribution function P(r), which was calculated from the entire scattering patterns via indirect Fourier inversion of the scatter-ing intensity I(q) using the program GNOM (42). The maximum particle diameter Dmax was also estimated from the P(r). The hydrated volume VP of the particle was computed using the Porod equation: VP = 2π2I(0)/Q, where I(0) is the extrapolated scattering intensity at zero angle and Q is the Porod invariant. (29,43). The molecular mass of a globular protein can then be estimated from the value of its hydrated volume following the method of Rambo and Tainer as implemented in the SAXNS_ES server [PMID:23619693] (29). The overall shape of the protein was modeled ab initio by fitting the SAXS data to the cal-culated SAXS profile of a chain-like ensemble of dummy residues in reciprocal space us-ing the program GASBOR, version 2.3i (44). Twenty-five independent calculations were performed with a D5 symmetry restriction (see below for choice of symmetry).

### Analytical Ultracentrifugation (**AUC**)

Analytical ultracentrifugation experiments were performed using an Optima XL-A analytical ultracentrifuge equipped with absorbance optics (Beckman Coulter, Brea, CA). Purified terminase ring was dialyzed into 20 mM Tris buffer, pH 8 at 4 °C containing 500 mM NaCl, 5% v/v glycerol and 1 mM TCEP and then concentrated using 10 kDa MWCO Amicon Centrifugal Filters (Millipore Sigma) according to manufacturer’s directions. The enzyme concentration was determined spectrophotometrically (ε280 = 179,680 cm^−1^ M^−1^ for the protomer) and then diluted using the dialysis buffer to the concentration indicated in each individual experiment. Buffer viscosity (1.893 mPa*s) and buffer density (1.03812 g/cm^3^) were measured at 4 °C using a Lovis 2000 M rolling ball viscometer/densitometer (Anton Paar). A 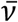 of 0.72364 mL/g at 4 °C and 0.73045 mL/g at 20 °C was determined using the Sedntrp program (Biomolecular Interactions Technology Center, University of New Hampshire, Durham, NH) and corrected for the change in hydration shell due to the presence of glycerol to yield 0.72397 mL/g at 4 °C and 0.73078 mL/g at 20 °C as described elsewhere.

For the *sedimentation velocity e*xperiments (**SV-AUC**), the samples (410 ml) were loaded into the sample cell of two sector Epon charcoal-filled centerpieces and dialysis buffer was loaded into the reference cell. The samples were allowed to equilibrate to 4°C and then spun at 32,000 rpm; sedimentation was monitored by absorbance at 250 nm. The data were analyzed using the SEDFIT program to afford c(s) distributions (45). The bottom fitting limit was moved to 6.7 cm to improve the fit accuracy by avoiding the glycerol gradient created at the cell bottom as reported elsewhere.

For the *sedimentation equilibrium* experiments (**SE-AUC**), 80 μL of each sample at the indicated concentration was loaded into the reference cell of six sector Epon-filled charcoal centerpieces and dialysis buffer was loaded into the reference cell. The samples were allowed to equilibrate to 4°C and then successively spun at 7,500, 9,000 and then 11,000 rpm. Sedimentation at each speed was assumed to be at equilibrium when consecutive scans, separated by intervals of 2 h, did not change as determined using the Heteroanalysis program (46). Absorbance data were collected at 250 nm every 0.003 cm in the step mode, with 5 averages per step. The protomer e250 (69373 M^-1^cm^-1^) was determined using a concentration curve and fit using a linear least squares approach (See Figure SX). A global non-linear, least-squares (**NLLS**) analysis of the combined SE data was performed using the SedAnal program (47) assuming a single-species model;

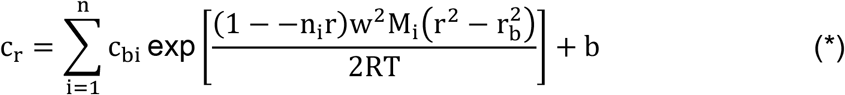

where c_bi_, v_i_, and M_i_ are the concentration at the bottom of the cell, partial specific volume and molecular mass of the “I” component, respectively; ρ is the density of the solution; ω is the angular velocity; and b is the base-line error term.

### Mass Spectrometry

The terminase samples were exchanged into ammonium acetate solution, a volatile salt used in native electrospray, by Zeba microbiospin columns, 7 kDa MWCO (Thermo Scientific). Terminase protomer was exchanged into a 200 mM ammonium acetate salt, and assembled ring was exchanged into a 300 mM ammonium acetate salt pH 8.34. After buffer exchange samples were immediately electrosprayed for CDMS analysis.

Charge detection mass spectrometry (**CDMS**) measurements were performed on a home-built instrument described in detail elsewhere (48-55). Briefly, the mass to charge ratio (*m/z)* and charge are simultaneously measured on thousands of individual particles and binned to give a mass distribution of the sample. Ions enter the instrument through a metal capillary after formation from a nanoelectrospray (nESI) source. The positively charged ions are guided through a series of ion optics and differential pumping to an electrostatic linear ion trap (ELIT) where they oscillate back and forth through a detection cylinder. Signal from the oscillating ion is picked up on a charge sensitive amplifier where it is digitized and processed using a fast Fourier transform. The fundamental frequency measured corresponds to the *m/z* of the ion and the magnitude corresponds to the charge.

### cryoEM Methods

A quantifoil R2/1 holey carbon copper grid (Electron Microscopy Sciences, Hatfield, PA, US) was plasma-cleaned for 40 s. A total of 3 μL of purified terminase at 0.85 mg/mL was applied to the grid prior to freezing using a Vitrobot automated plunge-freezing robot. Grids were loaded into a Krios Titan microscope housed at SCSB center at UTMB, and data collected at 300 KeV on a K3 direct electron detector operating in counting mode at 96K magnification at the detector, thus corresponding to a pixel size of 0.85 Å/pixel. *Each area was exposed to a total dose of 60 electrons per square Angstrom*. Individual movie frames were aligned and averaged using MotionCor2 (56) as implemented in RELION (57), with latter frames low-pass filtered to remove artefacts from radiation damage. CTF parameters for individual frames were determined and phases and amplitudes corrected for using the RELION implementation of Gctf (58). Particle picking and 2D classification were done using RELION.

### ATPase Kinetic Analysis

ATPase assays were performed by published procedure (34,59), with modification. Briefly, reaction mixtures (10 μl) contained 50 mM Tris-HCl, pH 9, 10 mM MgCl2, 60 mM NaCl, 2 mM spermidine, 7 mM β-ME, 5 μM [α^32^P] ATP, and 10 nM lambda terminase. Where indicated, DNA and IHF were added to a final concentration of 50 nM duplex (≈14 μM total nucleotide) and 60 nM protein, respectively. The reaction mixtures were incubated at 37°C for 5 minutes and were initiated with the addition of terminase and allowed to proceed for 20 minutes at 37°C. Aliquots (2 μl) were removed from the reaction mixture and quenched with the addition of equal volume stop solution (100 mM EDTA, 10 mM each cold ATP and ADP). Aliquots (2 μl) of the quenched reaction mixtures were spotted onto a silica gel TLC plate and the plate was developed with 60% of 2-propanol and 30% ammonium hydroxide. The images of ^32^P labeled ATP and ADP spots were captured by GE-Typhoon Phosphorimager and quantitated using the ImageQuant software. Control experiments (ATP hydrolysis time course) using these conditions were performed to ensure that the aliquots were taken within the linear portion of the reaction curve. The kinetic constants for ATP hydrolysis were determined by non-linear regression analysis of the experimental data using the Igor data analysis program (Wave Metrics, Lake Oswego, OR).

## Footnotes

1 Notable exceptions to this general paradigm are represented by the ϕ29-like bacteriophages and the adenovirus groups, which replicate monomeric genomes in a protein-primed manner and thus have no maturation requirement. Notwithstanding, these viruses utilize an analogous “packaging ATPase” enzyme to package viral DNA into a pre-assembled procapsid shell.

## Notes

### Competing Interest Statement

The authors have declared no competing interest.

